# Vertebrates show coordinated elevated expression of mitochondrial and nuclear genes after birth: a conserved metabolic switch

**DOI:** 10.1101/2024.06.25.600612

**Authors:** Hadar Medini, Dan Mishmar

## Abstract

Interactions between mitochondrial and nuclear factors are essential to life. Nevertheless, the importance of coordinated regulation of mitochondrial-nuclear gene expression (CMNGE) to changing physiological conditions is poorly understood, and is limited to certain tissues and certain organisms. We hypothesized that CMNGE is important for development across vertebrates, and hence will be conserved. As a first step, we analyzed >1400 RNA-seq experiments performed during embryo development, neonates and adults across vertebrate evolution. We found conserved sharp elevation after birth of CMNGE, including oxidative phosphorylation (OXPHOS) and mitochondrial ribosome genes, in heart, hindbrain, forebrain and kidney across mammals, *Gallus gallus* and in the lizard *Anolis carolinensis*. This was accompanied by elevated expression of TCA cycle enzymes, and reduction in hypoxia response genes, suggesting a conserved cross-tissue metabolic switch after birth/hatching. Analysis of ∼70 known regulators of mitochondrial gene expression revealed consistently elevated expression of PGC-1a and C/EBPB after birth/hatching across organisms and tissues, thus highlighting them as candidate regulators of CMNGE upon transition to the neonate. Analyses of *Danio rerio*, *Xenopus tropicalis* and *Drosophila melanogaster* revealed elevated CMNGE prior to hatching, coinciding with the development of motor neurons. Lack of such ancient pattern in mammals and in the chicken suggests that it was lost during radiation of terrestrial vertebrates. Taken together, our results suggest that regulated CMNGE during embryogenesis and after birth, alludes to metabolic switch which is under strong selective constraints and hence essential.

## Introduction

Mitochondria are pivotal to cellular metabolism across all eukaryotes. In animal cells, unlike most cellular pathways, the mitochondrial genetic system is the only system with factors encoded by two genomes, in two cellular compartments – the nucleus and the mitochondria. We and others previously found that coordination of gene expression between the mitochondrial (mtDNA) and nuclear genome, especially in the energy-producing oxidative phosphorylation system (OXPHOS) is common across most healthy human tissues (Barshad et al. 2018), a variety of cancer samples (Reznik et al. 2017), and in certain cell types (Medini et al. 2021a) yet is compromised in several disease conditions, including COVID19, Alzheimer disease and several cancer types (Medini et al. 2021b; Papier et al. 2022). It is thus reasonable to assume that coordinated mito-nuclear gene expression (CMNGE) is important, will respond to changing environments, and will change upon alteration of physiological conditions, such as the dramatic changes that occur during embryo development and in the transition to the neonate (Sharma et al. 2014).

Indeed, mitochondrial activity, morphology, number and regulation of replication and transcription are critical for cell differentiation and development (Folmes et al. 2011; Hom et al. 2011; Wellen and Thompson 2012; Medini et al. 2020). Specifically, knockout of mitochondrial regulatory proteins such as mtDNA polymerase gamma (POLG) (Hance et al. 2005) and mtDNA transcription factor A (TFAM)(Larsson et al. 1998) resulted in developmental arrest between embryonic days 7.5 and 8.5 (upon POLG knockout) and lethality on day 10.5 of mouse embryos (upon TFAM knockout). In addition, knockout of mitochondrial RNA polymerase (POLRMT) resulted in decreased brood size and impaired ova development in the worm *Caenorhabditis elegans* (Charmpilas and Tavernarakis 2020). Secondly, knockout of NRF1, a key regulator of nuclear DNA-encoded OXPHOS genes expression, led to mouse embryonic lethality (Chan et al. 1998). Third, knock out of another regulator of OXPHOS genes expression, NRF2, led to reduction in the levels of NADPH and the NADPH/NADP+ ratio in embryonic mouse fibroblasts (Singh et al. 2013) and ATP levels were significantly reduced upon Nrf2 knock down in human cancer cell lines (Kim et al. 2011). Notably, both NRF1 and NRF2 are involved in mitochondrial biogenesis via regulation of peroxisome proliferator-activated receptor-gamma coactivator-1alpha (PGC-1α) - an OXPHOS master regulator (Gureev et al. 2019a; Deng et al. 2020). Accordingly, targeted deactivation of PGC-1α as well as a transcription factor from the same family, PGC-1β (PGC-1αβ−/−) in the mouse heart led to reduced growth, late fetal arrest in mitochondrial biogenesis (at E16.5 and E17.5) and 70% lethality of mice within 24 h after birth (Lai et al. 2008). Fourth, dysfunctional mitochondria in the embryonic mouse and human patients’ heart lead to severe cardiomyopathy and embryonic or neonatal lethality (Zhao et al. 2019). Taken together, transcriptional regulation of mitochondrial genes in both the nucleus and the mtDNA is pivotal for the developing embryo and for the transition to the neonate.

Studies in model organisms support the occurrence of a metabolic shift after birth in certain tissues. Quantification of lactic acid in rat livers, revealed that glycolysis is decreased after birth (Burch et al. 1963). In consistence with this finding, human postnatal liver and heart collected 6 and 36 weeks after birth displayed an increase in mitochondrial content and respiratory chain activities as compared to fetal samples collected after 11–17 weeks of gestation (Minai et al. 2008). Finally, increased level of the mitochondrial electron carrier cytochrome c was observed in newborn human and rat livers as compared to fetal samples (Krizova et al. 2021).

These pieces of evidence led us to hypothesize that there is a regulated increase in OXPHOS function after birth, which is conserved in evolution. To test for this hypothesis, we interrogated N=1482 RNA-seq experiments from samples collected during embryo and fetal development as well as after birth from a variety of metazoans, in several different tissues.

## Results

### Postnatal mtDNA gene expression is generally elevated across evolution

To assess the dynamics of mitochondrial transcript regulation during metazoan embryo development we have analyzed publicly available bulk RNA-seq data from six mammals (*Homo sapiens* - human, *Macaca mulatta* - rhesus monkey, *Mus musculus* - mouse, *Rattus norvegicus* - rat, and *Oryctolagus cuniculus* - rabbit) and from *Gallus gallus* (chicken). The data was generated from seven tissues collected during embryogenesis and postnatal development (Dataset I, Table 1, and see ‘Resources’ in the Methods section) (Cardoso-Moreira et al. 2019; Chen et al. 2022). This analysis was augmented by analysis of RNA-seq from samples collected during *Sus scroffa* (pig) liver embryogenesis and from the neonate (Dataset II). To identify the specific time frame in which potential transition in gene expression occurred, the data was analyzed in sliding windows of available developmental stages, while testing a variable number of consecutive stages (e.g., windows containing either 2 or 4 of the tested developmental stages). Then, we compared the fold change of gene expression between subsequent stages within each window (Fig. 1B, and see Methods). Our findings revealed significantly elevated mtDNA gene expression, which was strikingly observed in postnatal samples from 5 out of 7 tissues across all tested species, with one exception – reduction in mtDNA gene expression in chicken liver after hatching (Fig. 1A, and see below). Specifically, we found elevated mtDNA genes’ expression right after or one week after birth/hatching in: heart in 6/6 of the tested organisms, forebrain/cerebrum in 5/6 organisms (human, rat, rabbit, mouse and rhesus monkey), hindbrain/cerebellum samples in 5/6 organisms (human, rat, mouse, rabbit and rhesus), kidney samples in 4/6 organisms (human, mouse, chicken and rhesus), liver samples in 5/7 organisms (human, rat, mouse, rabbit and pig – the only tissue tested in the latter), and in testes only in rhesus monkey (Table 2). Further support for the elevation of mtDNA gene expression after birth came from analysis of other additional RNA-seq dataset collected from postnatal mouse heart samples as compared to embryonic samples (Dataset IV - embryos collected from gestational day 13.5 versus samples collected 6 weeks after birth) (Matkovich et al. 2014) (Fig. S1). All these results reflect a conserved elevation of mito-nuclear gene expression across tissues, implying a conserved regulatory mechanism.

**Figure 1.**
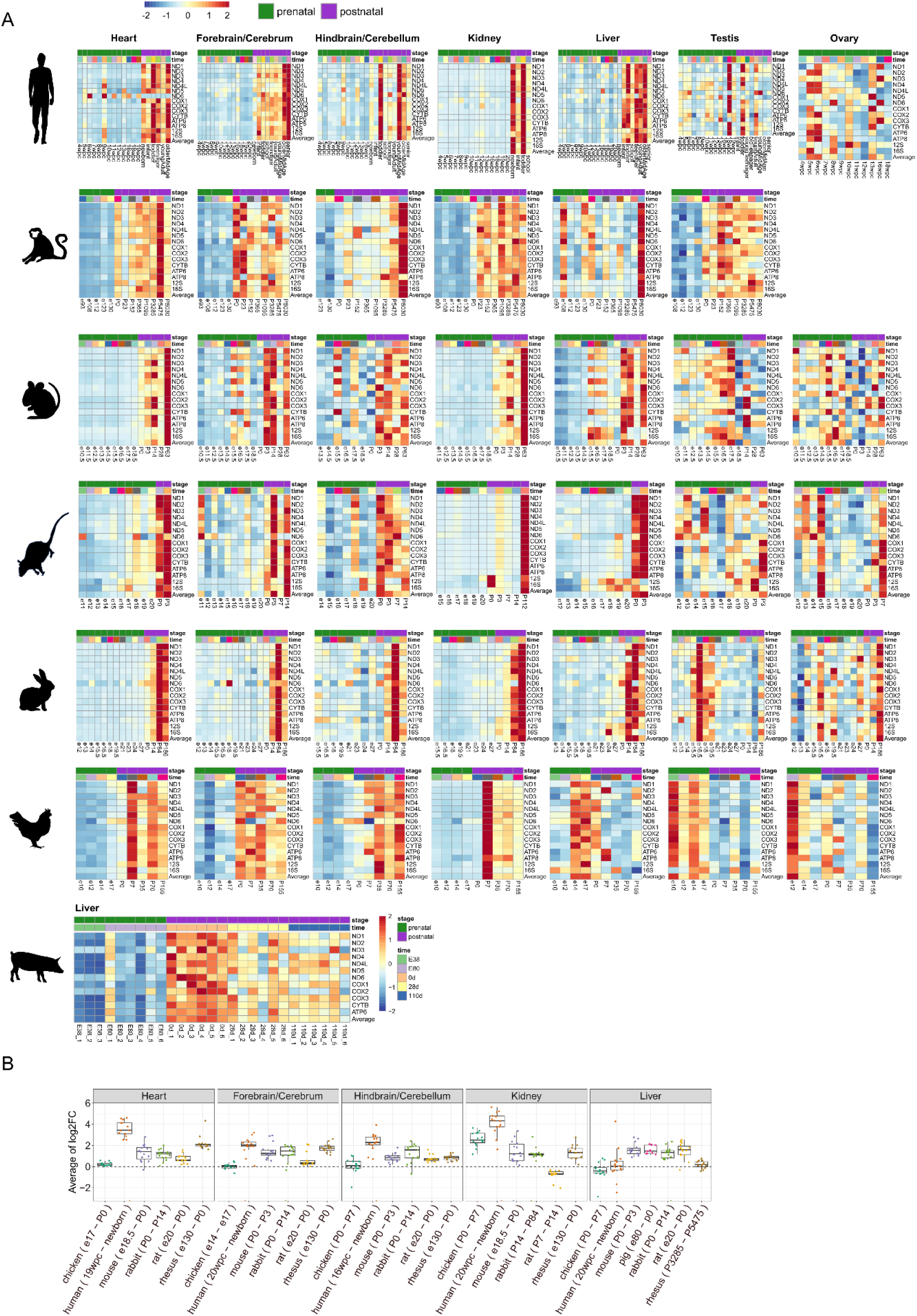
mtDNA genes expression during prenatal and postnatal development in vertebrates. (A) Heatmaps representing mtDNA gene expression in six tissues from five mammals and a non-mammalian vertebrate (chicken). Color bar indicates the scaled expression. Red scale: elevated expression, blue scale: reduced expression. X axis: prenatal (green) and postnatal (purple) stages of sample collection; we calculated the median expression level of samples from each given developmental stage. Y axis: mtDNA genes. (B) Boxplots representing log2FC values showing developmental stages with the highest FC in a window of four stages. X axis: developmental stages for each organism in different tissues.

**Table 1.** Resources for RNA-seq data per tissue and organism.

| Dataset | Organism/s analyzed | Tissue | Include P0 (neonate right after birth) | Resource |
| --- | --- | --- | --- | --- |
| I | Human (N=297), Rhesus (N=168), Mouse(N=316), Rabbit(N=315), Rat(N=350), Chicken(N=215) | Heart, Cerebrum/Hindbrain, Cerebellum/Forebrain, Kidney, Liver, Testis, Ovary | Yes | (Cardoso-Moreira et al. 2019) |
| II | Pig (N=30) | Liver | Yes | (Chen et al. 2022) |
| III | Chicken (N=9) | Liver | No | (Xu et al. 2019) |
| IV | Mouse (N=19) | Heart | No | (Matkovich et al. 2014) |
| V | *Anolis carolinensis* (N=78) | Brain, Heart, Kidney, Liver | No | (Marin et al. 2017) |
| VI | *Xenopus tropicalis* (N=40) | Whole embryo | No (only prenatal) | (Mitros et al. 2019) |
| VII | *Danio rerio* (Zebrafish) (N=56), *Drosophila melanogaster*(N=77) and *C. elegans* (N=81) | Whole embryo | No (only prenatal) | (Levin et al. 2016) |

**Table 2.** The developmental time point presenting the highest fold-change of either upregulation or downregulation of mtDNA gene expression in samples from Datasets I and II. Downregulation is indicated by (*) and the rest were upregulated. Numbers in parentheses indicate the level of coordination of nuclear DNA-encoded OXPHOS genes and mtDNA-encoded genes as explained in the Methods section.

|  | **Before P0** | **First week/s after birth** | **Right after birth/hatching** | **Late change** |
| --- | --- | --- | --- | --- |
| **Heart** | - | Rabbit(0.83) | Human(0.87),<br>mouse(0.88),<br>rat(0.72),<br>rhesus(0.86),<br>chicken(0.66) | - |
| **Forebrain/Cerebrum** | Chicken(0.35) | Mouse(0.37),<br>rabbit(0.59) | Human(0.72),<br>rat(0.06),<br>rhesus(0.65) | - |
| **Hindbrain/Cerebellum** | - | Mouse(0.69),<br>rabbit(0.52),<br>chicken(-0.19) | Human(0.62),<br>rat(0.19), rhesus<br>(0.41) | - |
| **Kidney** |  | Chicken(0.16),<br>rat(0) | Human(0.78),<br>mouse(0.84),<br>rhesus(0.88) | Rabbit(0.79) |
| **Liver** |  | Chicken(*)(-0.6),<br>mouse(0.67),<br>rabbit(0.27) | Human(0.23),<br>pig(0.8),<br>rat(0.03) | Rhesus(0) |
| **Testis** | Human,<br>rabbit, rat(*) |  | Chicken(*),<br>rhesus | Mouse(*) |
| **Ovary** | Chicken(*),<br>rabbit, rat(*) |  |  |  |

While considering the liver we noticed that unlike mammals, chicken liver displayed a reduction in mtDNA gene expression during the first week after hatching. This finding gained support from analysis of an additional dataset (Dataset III), which includes chicken liver samples from embryo developmental day 13 compared to samples collected 5 weeks after hatching (Fig. S2) (Xu et al. 2019). We interpret the divergent mtDNA gene expression pattern in the chicken liver as compared to mammalian liver as likely associating with the accumulation of fat in the chick liver, but not in mammals, thus offering a different carbon source for OXPHOS activity in chicken liver after hatching (see Discussion).

Our analysis so far focused mostly on mammals with one non-mammalian vertebrate – the chicken. To assess whether our observed elevated mtDNA gene expression after birth is shared across other vertebrates we sought to extend our analysis to other additional non-mammalian vertebrates. Accordingly, analysis of RNA-seq experiments generated from 48 embryonic samples from the lizard *Anolis carolinensis* (collected from developmental stages 15-16) as compared to 30 adult samples from four organs (brain, heart, kidney, liver) (Marin et al. 2017) (Dataset V, Fig. S3) revealed elevated mtDNA genes expression in the adult heart, brain, and kidney. Notably, the *Anolis carolinensis* liver did not show any consistent pattern (Fig S3). This finding suggests that the elevated mtDNA gene expression after birth is conserved across mammalian and terrestrial non-mammalian vertebrates.

### Postnatal nuclear DNA-encoded OXPHOS gene expression is elevated across evolution

As mtDNA-encoded proteins interact with nuclear DNA-encoded subunits of the OXPHOS system, we sought to assess the expression pattern of the latter during fetal development and after birth. Our findings indicate that the same tissues that showed elevated mtDNA gene expression also showed clear elevation of nuclear DNA-encoded OXPHOS gene expression in Dataset I, II, IV (Fig. 2A, 2B, Table 2, Fig. S4-S5). This finding supports the hypothesis that the elevation of OXPHOS genes’ expression after birth/hatching is coordinated between the mtDNA and the nucleus (mito-nuclear). To quantify the level of coordinated mito-nuclear gene expression in each of the tested organisms and tissues, we calculated the number of significantly altered expression of mtDNA genes and nuclear DNA-encoded OXPHOS genes and divided them by the number of genes with expression information (in each dataset, separately). This analysis revealed that the heart displayed high coordination level in all organisms analyzed (Average 0.8, SD 0.08) (Table 2, Fig. 2B, Fig. S4, Table S1). Moreover, the organisms that displayed change in mtDNA gene expression during the first weeks after birth/hatching also displayed coordinated mito-nuclear expression in the same days, including chicken liver that showed coordinated downregulation in OXPHOS genes (level of coordination = −0.6, Dataset I)(Fig. 2, Table 2, Table S1, Fig. S6). An exception was chicken hindbrain/cerebellum, that displayed elevated mtDNA gene expression between P0-P7 days after birth, yet decreased OXPHOS nuclear genes’ expression during the same days (level of coordination = −0.19). This observation awaits confirmation once additional RNA-seq datasets become available from chicken samples collected from the hindbrain during development. In *Anolis carolinensis* heart, and to a lower extent in the kidney, we observed a tendency towards elevated expression of structural nuclear-encoded OXPHOS genes (Fig S7). Notably, *Anolis carolinensis* brain did not show a consistently elevated OXPHOS gene expression in the adult. These results suggest that CMNGE is present in the heart and the kidney of the lizard *Anolis carolinensis*. Finally, it is worth noting that while considering mammals and chicken, structural subunits of the OXPHOS showed a sharper elevation in gene expression after birth than OXPHOS assembly subunits (Fig. 2A, 2B, Fig. S4). This suggests that while considering OXPHOS, elevated CMNGE after birth/hatching is more coordinated among the inherent structural members of the OXPHOS system.

**Figure 2.**
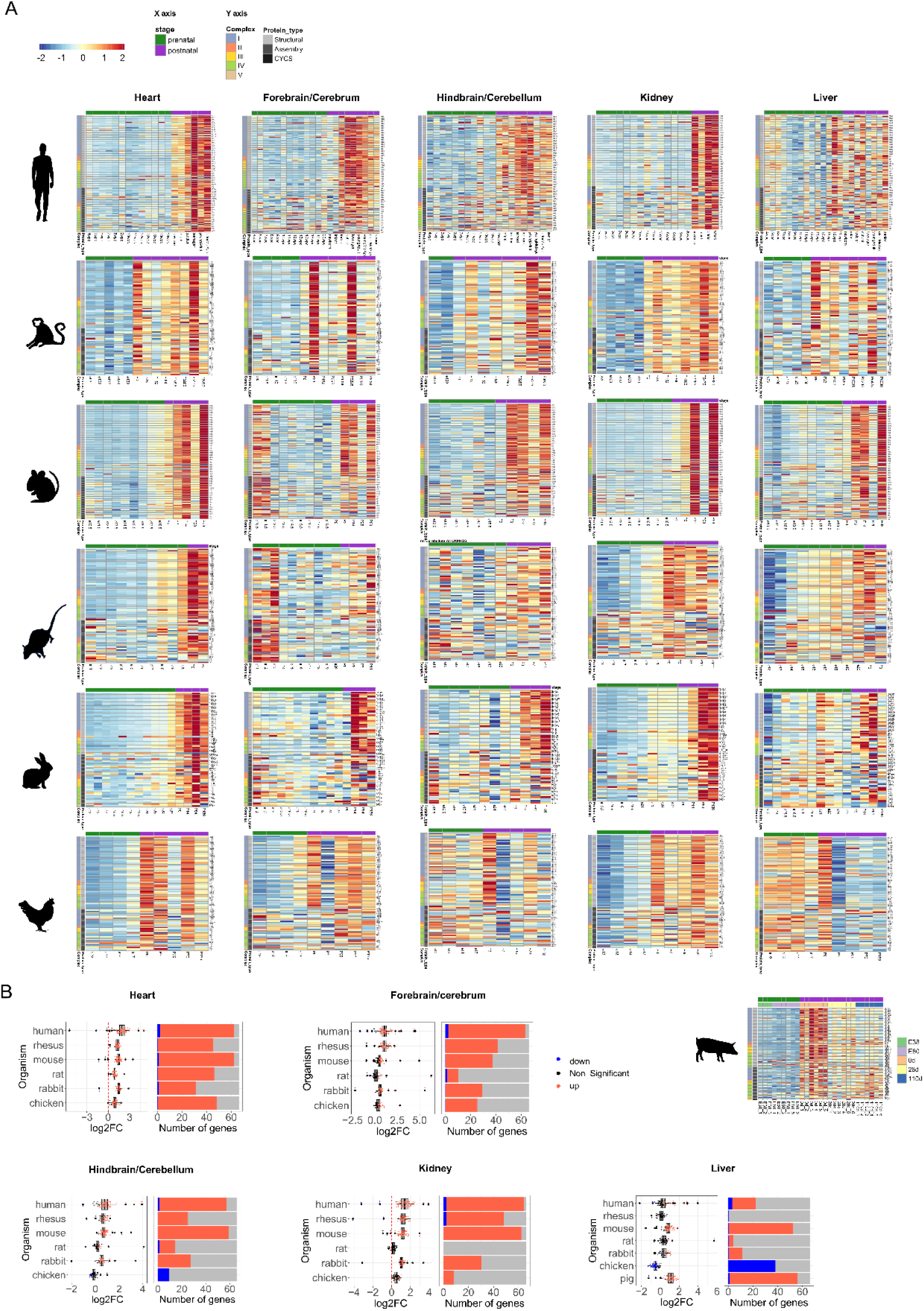
Nuclear DNA-encoded OXPHOS genes expression in embryo development and postnatal stages across vertebrates. (A) Heatmaps representing expression of nuclear DNA-encoded OXPHOS gene in six tissues from five mammals (six mammals in liver) and a non-mammalian vertebrate (chicken). Color bar: Red scale – elevated expression, blue scale: reduced expression. For each developmental stage the median expression of the relevant samples was used for calculation (Dataset I). Above each panel - horizontal green and purple bars span prenatal and postnatal samples, respectively. X axis: developmental stages. Y axis: nuclear DNA-encoded genes, divided according to OXPHOS complexes, structural and assembly genes and Cytochrome C (CYCS). (B) Left side of each panel: box plots indicating the log2FC of nuclear DNA-encoded structural factors of the OXPHOS. X axis: Log2FC. Y axis: the tested organisms. Color bar: blue dots-downregulation (log2FC< −0.2, p-value<0.05), red dots-upregulation (log2FC>0.2, p-value<0.05), black dots-non-significant. Dashed vertical red line displays the value of zero log2FC on x axis (no change). Right side of each panel: bar plots representing the number of significant (indicated in red or blue) and non-significant (NS) nuclear OXPHOS genes expression (colored in grey). Blue: statistically significant genes with decreased expression between the time points (p<0.05); Red: number of genes with significantly increased expression. X axis: Number of genes. Y axis: organisms.

### Altered mito-nuclear OXPHOS gene expression after birth is part of a wider metabolic switch

Since elevation in OXPHOS function after birth could be part of a general functional mitochondrial shift, we asked whether gene expression of any other mitochondrial pathways is altered after birth. Therefore, we compared gene expression of the following mitochondrial-related pathways before and after birth: the mitochondrial ribosome, the tricarboxylic acid (TCA) cycle, formation and scavenging of reactive oxygen species (ROS), in addition to glycolysis.

Firstly, the mitochondrial ribosome, which like the OXPHOS incorporate factors encoded by the nuclear and mitochondrial genomes – showed coordinated mito-nuclear expression change in heart of all organisms (Fig. S8, Table S1, Average 0.62, SD 0.24), and in the kidney of human, rhesus and rabbit (levels of coordination=0.64, 0.87 and 0.65, respectively). In addition, we observed strong CMNGE in the pig liver (Dataset II, level of coordination=0.78), to a lower extent in the mouse, but not in the liver of other tested mammals; interesting we observed reduction of gene expression in the chicken liver as in OXPHOS genes (level of coordination= −0.52). Finally, we found significantly decreased expression of the ribosomal genes in the cerebellum only in the chicken, similarly to the OXPHOS genes (level of coordination= −0.5) (Fig. S8). These results suggest that the mitochondrial ribosomal genes undergo a similar expression change after birth as in OXPHOS especially in the heart.

Secondly, we identified sharply elevated expression of genes related to the TCA cycle mainly in the heart and kidney after birth/hatching, and in several organisms in the rest of the tested tissues (Fig. S9). Notably, chicken liver displayed reduced TCA cycle genes’ expression, like in OXPHOS genes and the mitochondrial ribosome. Taken together, these results support a conserved overall functional metabolic shift in the mitochondria after birth or hatching across tissues and across the evolution of vertebrates.

Notably, our analysis of genes involved in glycolysis did not reveal any consistent overall expression pattern change during embryogenesis while jointly considered as a pathway (Fig. S10A). Nevertheless, lactate dehydrogenase (LDH), a key glycolysis enzyme, showed an interesting pattern: LDH has two subunits - LDHA and LDHB, with the first known to be upregulated mostly in glycolytic conditions, whereas LDHB is commonly upregulated in conditions that require OXPHOS (Read et al. 2001; Porporato et al. 2011). Although LDHA did not show any significantly conserved expression pattern across all tested tissues, it decreased in the heart after birth in 4/6 tested species (Fig. S10B, Fig. S10C). LDHB was significantly upregulated in several tissues that displayed mito-nuclear upregulation across species: the forebrain and hindbrain in 3/6 organisms, in the kidney in 4/6 organisms, and in the heart – in 5/6 of the tested organisms (Fig. S10B). Notably, the expression of enzymes and regulators of ROS scavenging pathway did not significantly alter in all tested mammals and in the chicken (Fig. S11). These results suggest that by and large, OXPHOS likely replaces glycolysis as the dominant ATP source after birth at least in the mentioned tissues. Nevertheless, the liver displayed significantly reduced LDHB expression after birth/hatching in 4/7 species (human, rhesus, mouse, rabbit, and chicken) (Fig. S10B, Fig. S10C). Since the liver is known to have special metabolic activities, including gluconeogenesis, we asked whether our observed unique expression pattern of the OXPHOS and LDHB associates with changes in the key enzyme of this pathway, Phosphoenolpyruvate carboxykinase (PEPCK) (Hanson and Reshef 1997). PEPCK (pck1, the cytosolic form) also has a mitochondrial isoform, pck2. Although PCK2 did not show any clear expression pattern change after birth/hatching, PCK1 was generally upregulated in all mammalian tissues and in chicken heart. Interestingly, the expression of PCK1 was significantly elevated in the liver of four mammals (human, mouse, rat and pig), yet did not significantly change in the chicken as well as in the chicken kidney after hatching – the main tissues where gluconeogenesis takes place (Fig. S12, Table S1). In summary, the generally elevated expression of PCK1 in the mammalian liver, lack of change in PCK1 in the chicken liver, along with only partial elevation of OXPHOS in this tissue in mammals, and reduction of LDHB expression in mammals, may reflect a unique involvement of the OXPHOS in liver metabolism during evolution.

*Anolis carolinensis* did not show any consistent pattern in these pathways while comparing embryo to adult samples (Figs. S13-S17, Table S1). This suggests that although the lizard did show a trend towards elevated OXPHOS gene expression in postnatal periods, more detailed expression analysis in this species after hatching may shed light on the involvement of other additional metabolic pathways.

Taken together, the CMNGE - coordinated upregulation of mito-nuclear gene expression (e.g., OXPHOS and the mitochondrial ribosome) is a widespread phenomenon across somatic tissues, is highly conserved among mammals and chicken, and is accompanied by significant elevation of TCA cycle enzymes. This supports an overall change in mitochondrial metabolism after birth/hatching which is regulated already at the transcript level.

### The elevated and coordinated mito-nuclear gene expression after birth associates with altered expression of regulatory factors that modulate mitochondrial biogenesis

The discovery of elevated CMNGE in vertebrates during the transition to the neonate suggests an underlying regulatory mechanism. Accordingly, it is conceivable that certain regulators of mito-nuclear transcripts modulate CMNGE elevation right after birth. To identify such factors, we tested for three categories: 1) transcription, 2) post-transcription 3) mitochondrial biogenesis. To this end we analyzed bulk RNA-seq samples from Datasets I and II including six mammals and chicken, for the expression pattern of ∼70 known regulatory factors of mitochondrial transcription, post-transcriptional activities, and biogenesis-related genes (Table S1, Table S2, Fig. 3, Figs. S18-S19). Specifically, we compared the expression of these genes in samples that precede the time point where the OXPHOS expression transition occurred to the samples right after the transition in each organism, per tissue.

**Figure 3.**
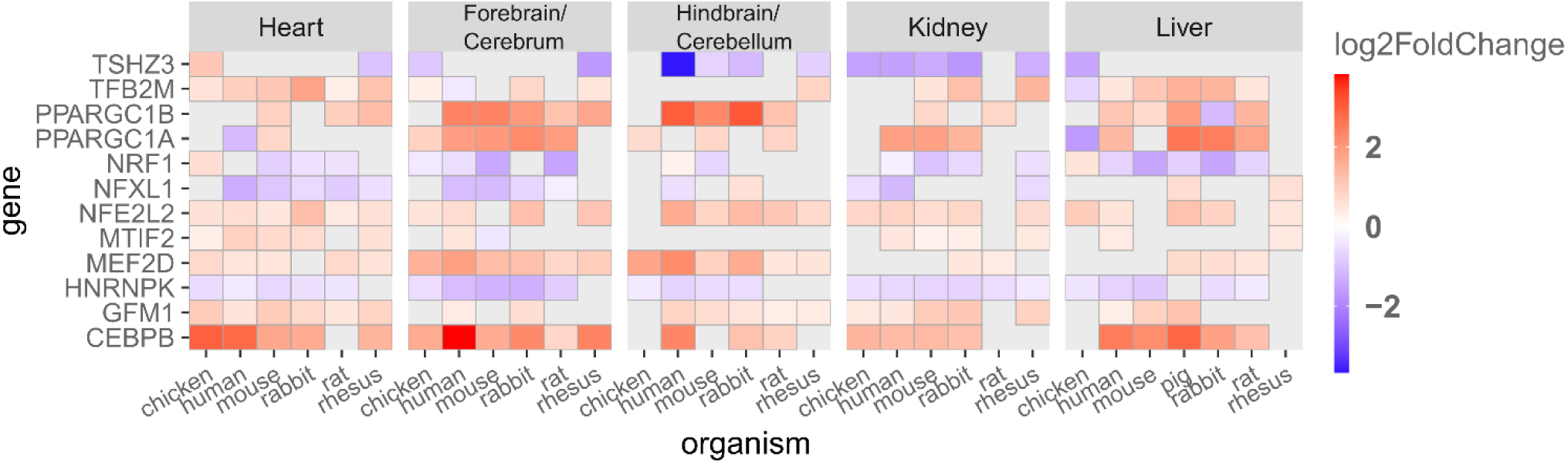
Expression of regulatory factors that modulate mitochondrial biogenesis associates with mito-nuclear expression shift after birth/hatching across vertebrates. Heatmap presenting log2FC values of regulatory factors and genes related to mitochondrial biogenesis that displayed significant change in at least four tissues within any organism and in at least five organisms within any tissue (p-value<0.05). Color bar indicates the log2FC. X axis: organisms divided by tissues, Y axis: genes. Only log2FC >0.2 and log2FC<-0.02 are shown.

As we expected that only a few factors will be involved in the elevated CMNGE in the neonate, we counted for each factor the number of organisms and corresponding tissues that displayed significant values in the tested factors. Specifically, we considered factors as candidates if they consistently displayed significantly altered expression in the transition to neonatal life in at least four tissues within any organism, and at least five organisms within any tissue (Fig. 3).

Firstly, our analysis highlighted Peroxisome proliferator-activated receptor gamma coactivator 1-alpha (PGC-1α), as a regulatory factor whose expression pattern best resembled the pattern of OXPHOS genes across organisms and tissues. PGC-1α is regulated by environmental stimuli and is considered central in regulating mitochondrial biogenesis (Puigserver et al. 1998). While considering the liver, the results revealed that PGC-1α was significantly upregulated in liver of 4/7 organisms: human (logFC=1.39), pig (logFC=2.64), rabbit (logFC=2.5) and rat (logFC=1.77) while in contrast it was downregulated in chicken liver (logFC= −1.65), in similar to mito-nuclear OXPHOS genes. The elevated expression of PGC-1α was also apparent in other tissues in the tested organisms (forebrain/cerebrum, hindbrain/cerebellum, and kidney), yet intriguingly was not apparent in the heart (Fig 3, Fig. S18). This result supports the association of elevated CMNGE after birth with altered regulation of mitochondrial biogenesis in most tested tissues.

Secondly, NFE2L2 which encodes Nuclear Respiratory Factor 2 (NRF2), a transcription factor that responds to oxidative stress and is involved in regulation of mitochondrial biogenesis via regulation of PGC-1α (Gureev et al. 2019a; Deng et al. 2020), was significantly upregulated in the following tissues and organisms (Fig. 3, Fig S18, Table S1): in the heart (all organisms), in forebrain/cerebrum (4/6 organisms), in hindbrain/cerebellum and the kidney of 5/6 organisms and liver of 5/7 organisms. Notably, in different from PGC-1α, NFE2L2 expression was elevated in chicken liver.

Third, CCAAT Enhancer Binding Protein beta (C/EBPB) was significantly upregulated in the heart of 5/6 organisms, in the forebrain/cerebrum of all organisms, in hindbrain/cerebellum of 3/6 organisms, in kidney of 4/6 organisms and in liver of 5/7 organisms (Fig. 3, Fig S18). Previous analysis of ChIP-seq experiments from the ENCODE consortium revealed that C/EBPB binds upstream regulatory elements of OXPHOS genes more than expected by chance (Blumberg et al. 2014a). This is the only transcription factor in the current analysis that was experimentally shown to both regulate nuclear gene transcription and bind the mtDNA *in vivo* (Blumberg et al. 2014a). Notably, C/EBPB is a transcription factor involved in inflammation, metabolism, and differentiation as well as in fatty acid metabolism in collaboration with PGC-1α (Takagi et al. 2022). Therefore, C/EBPB forms an attractive candidate to regulate transcription of both nuclear and mtDNA OXPHOS genes in the transition to the neonate.

Fourth, GFM1 (GTP dependent translation elongation factor mitochondrial 1), was upregulated in the heart (5/6 organisms), in the forebrain/cerebrum of 2/6 organisms, in the hindbrain/cerebellum of 5/6 organisms, in the kidney of 5/6 organisms, and in the liver of 3/7 organisms (Fig. 3). The upregulation of GFM1 is of interest in the context of upregulated mitochondrial ribosomal proteins in the neonate. This raises the possibility that mitochondrial protein synthesis is also upregulated in the neonate.

Notably, the expression of NRF-1, a known regulator of OXPHOS gene expression in the nucleus and of mitochondrial biogenesis (Gureev et al. 2019b), was significantly downregulated in the liver of 5/7 organisms (human, mouse, pig, rabbit, and rat) but elevated in chicken liver, and decreased in the kidney and forebrain/cerebrum of 4/7 organisms (Fig 3, Fig S18). This suggests that only a subset of the factors that regulate mitochondrial biogenesis form candidate regulatory factors to be involved in CMNGE after birth/hatching.

Finally, while considering the core mtDNA transcriptional regulatory factors we noticed that TFB2M was significantly upregulated in heart of all species, in forebrain/cerebrum of 3/6 organisms (chicken, rabbit and rhesus), in the hindbrain/cerebellum of rhesus, in the kidney of 3/6 organisms (mouse, rabbit and rhesus), and in the liver of 5/7 organisms (human, mouse, pig, rabbit and rat) (Fig. 3, Table S1). Notably, it was significantly downregulated in chicken liver. Then, while considering mitochondrial transcription factor A (TFAM) we found that it was significantly downregulated after birth in human heart (logFC=-0.63), forebrain/cerebrum samples from human, rat and rabbit (logFC=-1.37, logFC=-1.12, logFC=-0.81, respectively), hindbrain/cerebellum of human and rabbit (logFC=-0.81, logFC=-0.51, respectively), human and chicken kidney (logFC=-0.5, logFC=-0.34, respectively) and chicken liver (logFC=-0.61, respectively) (Fig. S18, Table S1). As in high cellular concentrations TFAM coats the mtDNA, and in lower concentrations, it promotes mtDNA transcription and replication, it is conceivable that lower TFAM levels reduce mtDNA occupancy and increase mtDNA transcription and replication (Kukat et al. 2015). In consistence with this interpretation, POLRMT was significantly upregulated in heart of human, mouse and rat, and human kidney; these findings support the possibility that core mtDNA transcriptional machinery partially participates in the CMNGE upregulation after birth (Fig. S18).

For *Anolis carolinensis* two genes that were also shared with mammals and chicken showed a significant elevation in expression: GFM1 in kidney (log2FC = 1.52) and C/EBPB in liver (log2FC= 2.81) (Fig. S20, Table S1).

Taken together, regulation of mitochondrial biogenesis seems the most prominent regulatory mechanism which responds similarly to the CMNGE shift in the neonate. Hence, it is conceivable that control of mitochondrial biogenesis is the main player that steers the mitochondrial-metabolic transition from fetus to neonate in mammals and in chicken.

### Hypoxia and locomotion: assessment of altered pathways which likely require the upregulation of CMNGE after birth

Mammalian embryos develop *in utero* in hypoxic conditions (Webster and Abela 2007). Previous work revealed that animal exposure to hypoxia reduces mitochondrial metabolism and allow cardiomyocyte regeneration in the neonate and in adult mouse heart (Nakada et al. 2017). Thus, we reasoned, that upregulation of CMNGE after birth mechanistically associates with the transition to an oxygen-rich environment and is important for the transition of embryonic to adult tissue. Therefore, as a first step, we tested for the expression of the top genes that are known to be upregulated in hypoxia in humans during development. Our results revealed a reduction in the expression of most of the genes that positively respond to hypoxia across tissues and species (Fig. 4, Fig. S21). We also noticed that humans display stronger reduction in brain and liver tissues compared to the heart and kidney. Secondly, in human heart ∼40% of the significant genes related to hypoxia were upregulated (Fig. S21).

**Figure 4.**
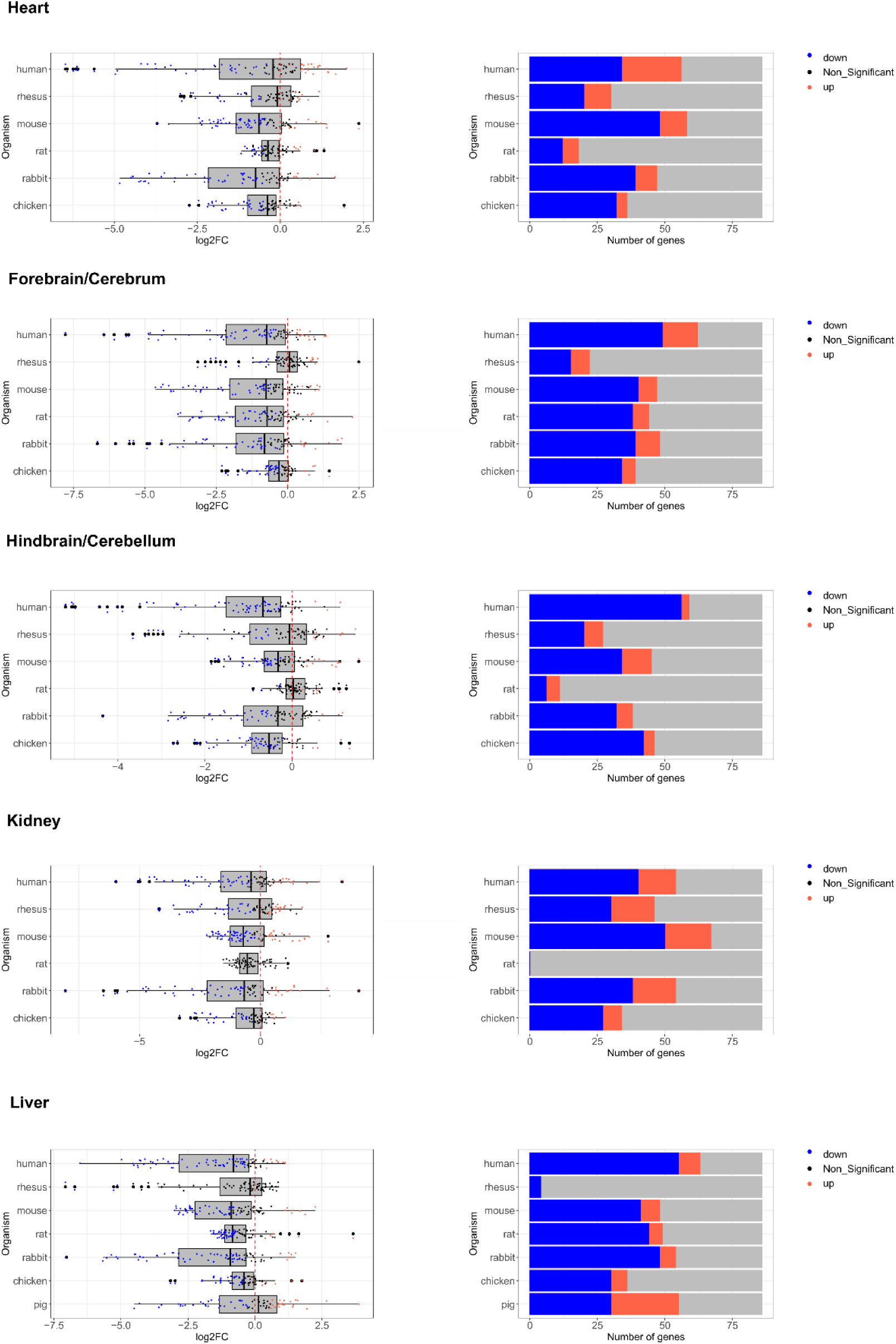
Hypoxia response gene expression is generally reduced after birth. <u>Left side panels</u>: box plots representing the log2FC of hypoxia genes. X axis: log2FC values, Y axis: organisms. Color bar: blue dots - significant downregulation (log2FC< −0.2, p-value<0.05); red dots - significant upregulation (log2FC>0.2, p-value<0.05); black dots - non-significant. Dashed vertical red line corresponds to the value of zero log2FC on X axis (no change). <u>Right side panels</u>: bar plots representing the number of significant (upregulated in red, downregulated in blue) or insignificant genes (in grey). X axis: number of genes, Y axis: organisms tested.

Since the transition to the neonate associates with exit from a confined space (the womb, egg) to the external environment, we hypothesized that expression of genes involved in regulation of locomotion will be altered. While analyzing genes related to locomotion, we did not find a consistent overall change (Fig. S22). Since the forebrain harbors most neurons that direct locomotion (Ferreira-Pinto et al. 2018) our findings suggest that the upregulation of CMNGE in the forebrain may contribute to the emergence of locomotion. As the activity of proteins involved in locomotion is logically related to skeletal muscle, it would be of interest to study the expression of this pathway in muscle after birth when such data becomes available.

*Anolis carolinensis* did not show any consistent pattern in these pathways while comparing embryo to adult samples (Figs. S13, S23-S24)

Together, the transition of the fetus from hypoxic conditions to breathing atmospheric oxygen, might influence the OXPHOS gene expression after birth, while a shift in the mode of locomotion, might not influence the OXPHOS gene expression in these tissues and organisms.

### Elevation of mito-nuclear gene expression before hatching associated with the development of locomotion neurons in *Xenopus tropicalis*, *Danio rerio* and *Drosophila melanogaster*

Although mammals and chicken displayed elevated CMNGE after birth, they did not show any clear CMNGE alteration before birth. We therefore asked whether putative changes in CMNGE during embryogenesis were masked by the major impact of postnatal samples. To address this possibility, we analyzed the same species focusing only on prenatal samples before birth/hatching. This analysis did not reveal any consistent alteration in CMNGE among the tested species (mammals and chicken) during embryo development (Fig. S25). We next analyzed bulk RNA samples collected during embryonic and fetal development from two non-mammalian vertebrates – *Xenopus tropicalis* (Mitros et al. 2019) (tropical frog, with two experimental repeats) and *Danio rerio* (Zebrafish) (Levin et al. 2016), as well as from two invertebrates - *Drosophila melanogaster* and *Caenorhabditis elegans* (*C. elegans*) (Levin et al. 2016) (whole organism). *Xenopus tropicalis* samples (whole organism) were collected from embryo cleavage stages until the tailbud free-swimming stage (Mitros et al. 2019), while *Danio rerio*, drosophila and *C. elegans* were sampled from the cleavage stage until hatching (but not after). While analyzing mtDNA and nuclear DNA-encoded OXPHOS gene expression, we found that unlike mammals and chicken, CMNGE elevation occurred in drosophila, zebrafish and in the frog before hatching (Fig. 5A). Notably, analysis of *C. elegans* revealed elevation in mtDNA gene expression at cleavage, yet it was accompanied by expression elevation of only several nuclear DNA-encoded genes from complexes I, III and IV towards hatching (Fig. 5A). These results suggest that the worm displays a much weaker CMNGE during development as compared to the other tested species.

**Figure 5.**
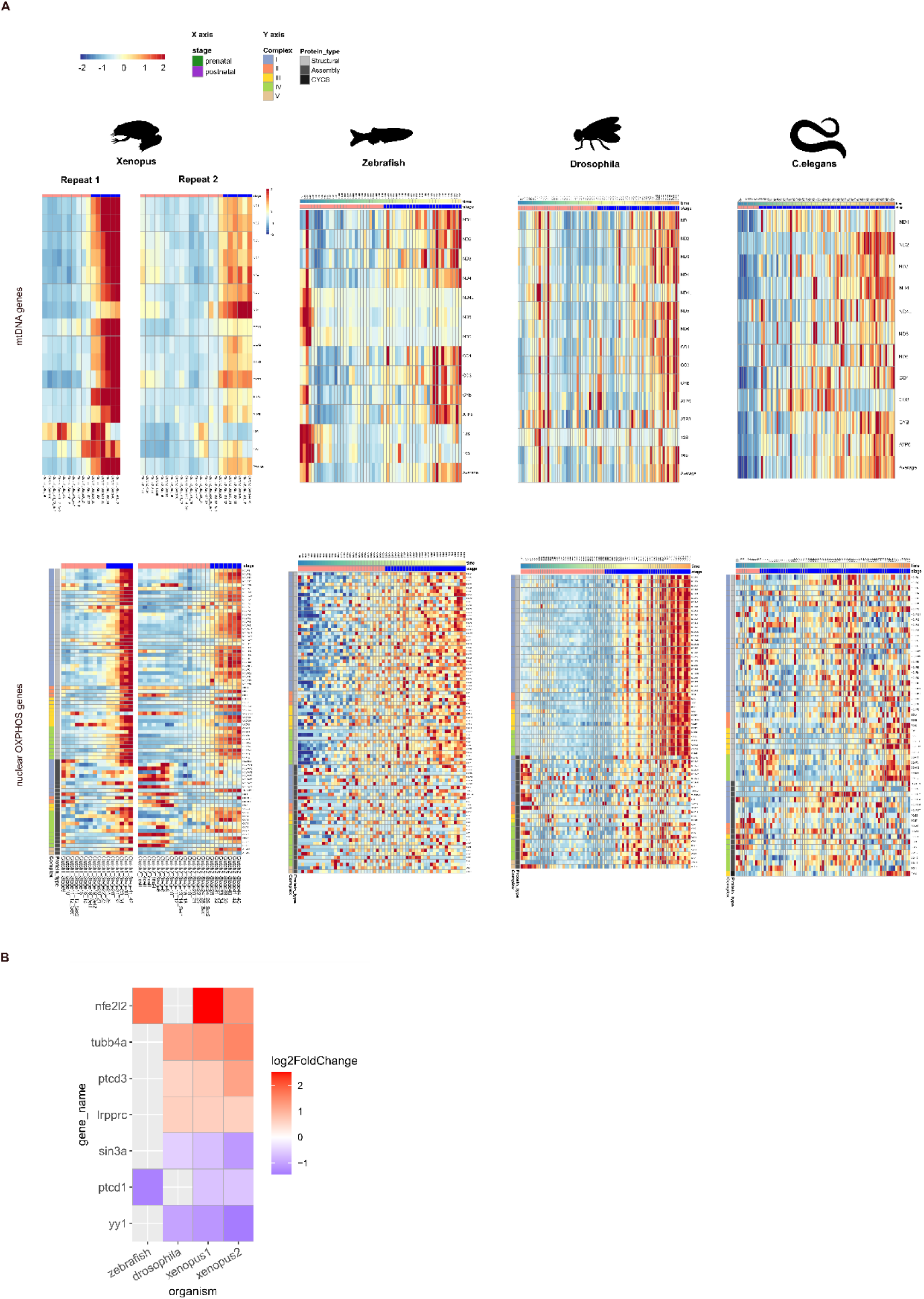
Mito-nuclear gene expression is elevated during embryogenesis. (A) Heatmaps of mtDNA and nuclear DNA-encoded OXPHOS gene expression from *Xenopus tropicalis*, *C.elegans*, *Drosophila melanogaster* and *Danio rerio*. Color bar indicates the scaled expression. Red scale: elevated expression, blue scale: reduced expression. X axis: prenatal samples, Y axis: mtDNA/OXPHOS genes. Notice division into OXPHOS complexes and into structural and assembly genes of the OXPHOS. (B) Heatmap representing log2FC values of regulatory factors and genes related to mitochondrial biogenesis. Color bar indicates the log2FC. X axis: organisms; Y axis: genes. Only log2FC >0.2 and log2FC<-0.02 are shown.

Our analyses raised the possibility that elevated mtDNA gene expression during development may display different timing while considering additional organisms – vertebrates and invertebrates (see below). Notably, while considering *Xenopus tropicalis* we found that CMNGE was increased during stages 24-26 of the first experimental replicate and at stage 28 of the second replicate, e.g. before hatching (Fig. 5A). Then, *Drosophila melanogaster* displayed elevation in CMNGE on 960 minutes, *Danio rerio* on 2120 minutes after fertilization. Interestingly, these stages in the frog and in drosophila coincide with initial motor neuron developmental and with the beginning of spontaneous movement (Fisher et al. 2023); in zebrafish the elevation in CMNGE occurred right after spontaneous movement commenced, yet in a more gradual manner (Fig. 5). Therefore, we analyzed genes related to the Locomotion pathway (Fig. S26A) and identified expression elevation in genes that were shared by at least two species while considering the frog, the fly, and the zebrafish (Fig. S26B). This analysis suggests that in these three species (frog, zebrafish, and drosophila) the timing of elevated CMNGE associates with the beginning of spontaneous locomotion, tempting us to speculate a causal connection.

We next asked whether the elevation of CMNGE towards hatching is unique to OXPHOS or could be found in other mitochondrial pathways of metabolism. Our analysis revealed that the expression of TCA cycle enzymes, and the mitochondrial ribosome genes were also elevated towards hatching in *Xenopus tropicalis,* Zebrafish and *Drosophila melanogaster* (Fig. S27). Notably, glycolysis did not show any altered expression pattern towards hatching.

Next, we asked whether CMNGE that was observed toward hatching in *Drosophila melanogaster* and *Xenopus tropicalis* is modulated by mitochondrial transcription factors, RNA binding factors or by biogenesis regulatory factors. The results highlighted three candidate regulatory genes that were shared between *Drosophila melanogaster* and *Xenopus tropicalis* (in two independent datasets): tubb4a (regulator of mitochondrial translation elongation), ptcd3 (binds mtDNA rRNAs and regulator of mitochondrial translation) and lrpprc (regulator of mitochondrial RNA stability). YY1, a known regulator of OXPHOS nuclear gene expression (Nandi et al. 2020), showed reduced expression in *Drosophila melanogaster* and *Xenopus tropicalis*, thus excluding it from positive involvement in regulating the elevated CMNGE during development in these species. An additional gene that displayed reduced expression in these organisms is sin3a, a transcriptional repressor protein known to regulate mitochondrial components. A previous study showed that loss of SIN3 in *Drosophila melanogaster* cultured cells results in up-regulation of both nuclear-encoded and mtDNA-encoded genes (Barnes et al. 2010). We noticed that only two genes displayed a shared expression pattern during embryogenesis of *Danio rerio* and *Xenopus tropicalis* – the first is nfe2l2 (NRF2) that showed elevated expression in similar to mammals after birth, and ptcd1 – that displayed reduced expression (Fig. 5B). Hence, *Drosophila melanogaster* and *Xenopus tropicalis*, but not *Danio rerio*, likely share the same mechanism controlling CMNGE during embryogenesis, which highlights post-transcriptional regulation and the regulation of translation as the most prominent underlying mechanism.

## Discussion

In the current study we assessed the cross talk between embryo development and regulation of mitochondrial function at the RNA level during metazoan evolution across multiple tissues (Fig. 6). To this end we analyzed transcriptomes that were collected during embryo development, as well as from neonatal and later stages from a variety of metazoans across several tissues. Our analysis showed concomitant increase of both mtDNA and nuclear DNA-encoded OXPHOS gene expression either immediately or shortly after birth (mammals) or either hatching (chicken) or postnatal periods (*Anolis carolinensis*). It was previously shown that the transition from the fetus to the neonate in the mouse heart requires a switch of postnatal metabolism to OXPHOS (Rashid et al. 2018; Zhao et al. 2019; Secco and Giacca 2023), and a profound alterations of mitochondrial protein functions in the early postnatal period (Talman et al. 2018). Our findings argue for a general, cross tissue, and evolutionarily conserved elevation in mitochondrial function after birth; we propose that these results imply that the transition of the embryo to the neonate most probably relies on CMNGE. Notably, while considering OXPHOS, most of the elevated CMNGE was observed in structural members of the OXPHOS rather than in the assembly genes. This suggests that CMNGE involves the former, implying that OXPHOS assembly genes are regulated differently from the structural OXPHOS genes during development. Secondly, along with the elevated mito-nuclear gene expression after birth we observed elevated expression of major mitochondrial metabolic pathways, such as the mitochondrial ribosome and TCA cycle enzymes. Alongside, and consistent with this finding, it was reported that the mtDNA copy number is significantly increased in human liver and muscle, supporting a general increase in mitochondrial mass after birth (Pejznochova et al. 2010). As we did not observe clearly consistent change in the expression of mtDNA replication factors after birth (Table S1), the reported change in mtDNA copy number might only reflect change in mitochondrial mass rather than mtDNA replication. Finally, we noticed that the coordinated mito-nuclear gene expression alteration after birth/hatching was evident in somatic tissues, but less in the reproductive organs (testis and ovary), apart from certain organisms. Since reproductive organs become more active upon puberty it would be of interest to assess mitochondrial gene expression in reproductive organs before and after puberty. Additionally, since mito-nuclear gene expression is likely coordinated also at the level of single cells (Medini et al. 2021a), it would be of interest to assess the mito-nuclear gene expression response in different cell types and to perform scRNA-seq analysis before and after birth from different tissues, hopefully from a variety of organisms. Taken together, our results strongly support a regulatory basis underlying the coordinated response of mitochondrial and nuclear OXPHOS gene expression at the RNA level to the transition from fetal to neonatal life.

**Figure 6.**
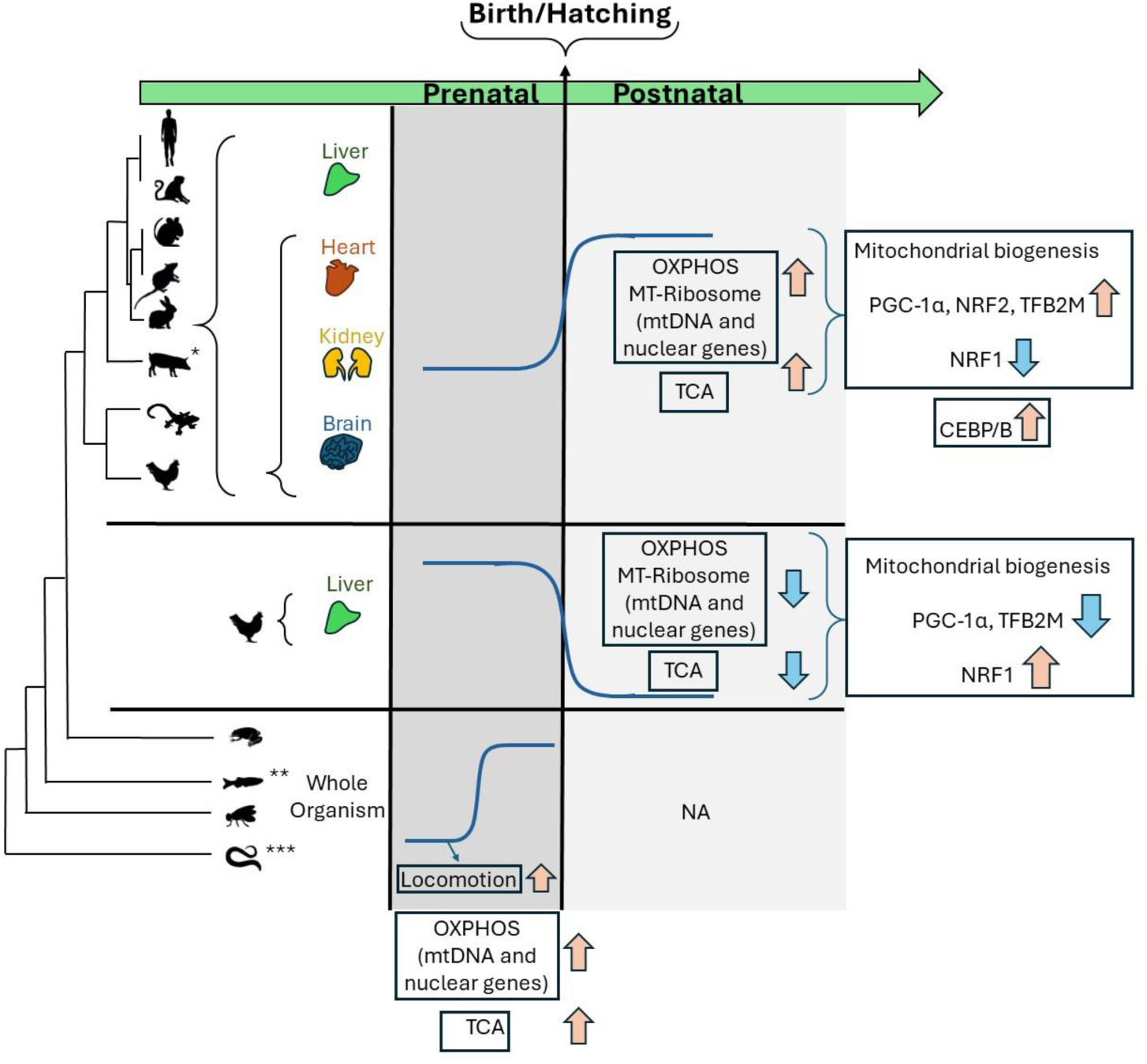
The landscape of mitochondrial-nuclear (mito-nuclear) gene expression during the course of development across metazoan evolution. Oxidative phosphorylation, mitochondrial ribosome, and TCA pathways showed elevated expression right after birth in mammals and in chicken across tissues. This pattern is associated with elevated expression of regulators of mitochondrial biogenesis and with hypoxia. Zebrafish, frog and drosophila showed coordinated mito-nuclear gene expression elevation in association with start of locomotion in the embryos. Blue arrows-downregulation, orange arrows-upregulation, the grey areas highlight the elevated or reduced mitochondrial gene expression. *-Sus scroffa (pig) included only liver samples. **-Danio rerio (Zebrafish) showed gradual elevation of mitochondrial genes. ***-C. elegans showed elevation of mtDNA gene expression similar to Danio rerio yet showed weaker coordinated expression pattern with nuclear mitochondrial genes.

One clear exception from the coordinated upregulation is the chicken liver: unlike the elevated CMNGE in mammalian liver samples, the chicken liver showed reduced mito-nuclear gene expression. After hatching, the chick’s main source of energy is derived from the conversion of feed-based carbohydrates into fatty acids, which mainly occur in the liver – as in humans (Noble and Cocchi 1990; Hicks et al. 2022). However, unlike humans and other mammals which harbor adipocytes in a variety of body sites, much of the fat of the chick is stored in the liver (Leveille 1969). It is thus conceivable that the hepatic molecular metabolic response to post-hatching in the chicken involves an increase in expression of genes involved in lipolysis (Latour et al. 1995). This is different from the mammalian neonatal liver which performs uptake of fatty acids from maternal milk, esterification of fatty acids into triglycerides, and storage in lipid droplets. Only upon breast feeding the neonatal liver may also engage in lipolysis and fatty acid oxidation as needed for energy production. These differences may impact the usage, and underlying striking regulatory difference of mito-nuclear gene-expression in the chick versus neonatal liver upon birth/hatching. Regardless, mito-nuclear gene expression is coordinately increased (mammals) or reduced (chicken), again supporting an underlying regulatory mechanism.

It is worth noting that although in general *Anolis Carolinensis* displayed elevated mito-nuclear gene expression in the comparison of embryos to postnatal samples, no such elevation was observed in other tested mitochondrial-related pathways. It is possible that more detailed analysis of lizard samples across embryo development and after hatching may unmask potential differences. This calls for detailed analysis of lizard samples right after hatching, like in the chicken. Such future analysis will be able to assess the conservation of our observed mitochondrial metabolic switch in the transition to the neonate.

To identify the best candidate factors to regulate the altered CMNGE after birth/hatching we screened a compendium of known regulators of transcription, post-transcription and mitochondrial biogenesis which displayed expression changes after birth across tissues and across evolution. Our findings highlighted an elevation of PGC-1α expression right after birth in many tissues and most organisms, except for chicken liver in which we observed decrease in PGC-1α expression – the very same pattern observed for mito-nuclear gene expression. With this in mind, we also found that the expression pattern of NRF2 – a known transcriptional regulator of nuclear DNA-encoded OXPHOS genes, which regulates PGC-1α transcription and numerous respiratory chain genes, including cytochrome c oxidase (COX) components

(Scarpulla 2002; Gureev et al. 2019a), also followed the mito-nuclear gene expression pattern across tissues and tested organisms. Interestingly, whereas NRF2 expression was elevated in the heart of most tested organisms, PGC-1α expression did not, suggesting that the upregulation of NRF2 in the heart after birth might be regulated by other factors. This interpretation should be tested in the future.

PGC-1α, a major regulator of mitochondrial biogenesis, acts as transcriptional activator not by direct DNA binding, but rather via protein-protein interactions with DNA-bound transcription factors (Tavares et al. 2020). Previous study showed that deactivation of PGC-1α in mouse hearts led to reduced growth, halted mitochondrial development late in gestation, and high lethality shortly after birth (Lai et al. 2008). As current data supports the limitation of PGC-1α activity to the nucleus, it is plausible that its involvement in regulation of mito-nuclear gene expression occurs via regulating the regulators of both mitochondrial and nuclear OXPHOS genes. This thought is partially supported by our observed elevated expression of the mitochondrial RNA polymerase, POLRMT, which is regulated indirectly by PGC-1α in mitochondrial biogenesis pathway (Wu et al. 1999; Gureev et al. 2019b), in tissues that showed significantly altered CMNGE in certain species. Accordingly, the expression of TFB2M, a transcription initiation factor that interacts with POLRMT (Falkenberg et al. 2002), was elevated after birth/hatching and its transcription is regulated by NRF-2 (Gleyzer et al. 2005). However, it was previously shown that a major regulator of mtDNA transcription and replication – transcription factor A (TFAM), is positively regulated by PGC-1α, as knockout of the latter caused reduced expression of the former (Gabrielson et al. 2014). Since our findings showed that TFAM expression is generally reduced after birth, the regulatory role of PGC-1α after birth is likely more limited. This interpretation is further supported by the following: Although the expression of PGC-1α is known to increase the expression of both NRF1 and NRF2 mRNAs (Wu et al. 1999; Gureev et al. 2019b; Popov 2020), we found that the expression of NRF1 was decreased after birth. It was previously shown that NRF1 positively regulates the activation of TFAM (49). Therefore, the elevated CMNGE is associated only with a subset of the regulators of mitochondrial biogenesis (see below).

The expression of C/EBPB was consistently upregulated after birth in most tissues and organisms tested, apart from the chicken liver. Unlike PGC-1α, C/EBPB has been *in vivo* localized both to the nucleus and to the mitochondria in cells, thus serving as an attractive candidate to directly participate in coordinated mito-nuclear gene expression. This interpretation awaits support from future silencing and overexpression experiments of C/EBPB in cells, which will enable to directly assess the impact of C/EBPB on CMNGE.

Since CMNGE after birth and hatching is common to all mammals, chicken and the lizard *Anolis carolinensis* we reasoned that there should be a common condition in the transition to the neonate, or neonatal phenotype, that most probably require such a sharp and conserved shift in mitochondrial function. The first condition to be considered is the transition from hypoxia to breathing atmospheric oxygen. With this in mind, it is important to mention that hypoxia suppressed the activity of PGC-1α and reduced its mRNA levels in cell lines (LaGory et al. 2015). Furthermore, hypoxia inducible factor (HIF) is essential for mediating PGC-1α suppression in hypoxia (LaGory et al. 2015), which is consistent with upregulation of PGC-1α upon birth. However, this explanation is likely tissue-specific in some organisms, as we found tissue dependent response of hypoxia regulatory factors after birth: although, as expected, most of the hypoxia genes were downregulated in human brain, liver and kidney, human heart displayed weaker effect in the expression of hypoxia responsive genes after birth. As the heart experiences a sudden elevation in oxygen level after birth one would expect a hypoxic response in the heart after birth due to oxidative stress (Secco and Giacca 2023) and in response to hypoxia – an elevated mitochondrial energy production in cardiac cells (Essop 2007). Therefore, despite the weaker reduction in the hypoxia pathway in human heart, our findings suggest that hypoxia may by-and large explain the need for elevated CMNGE after birth.

Although we did not identify a clearly consistent change in mito-nuclear gene expression during embryo development of mammals and chicken across tissues, we asked whether such changes could be identified in more distant metazoans including vertebrates and invertebrates. Indeed, our analysis of *Xenopus tropicalis*, *Danio rerio* and *Drosophila melanogaster* revealed significantly elevated CMNGE during the developmental stages when spontaneous embryo locomotion emerges. Specifically, the elevated CMNGE in *Xenopus tropicalis* coincides with the developmental stages in which initial motor neuron and spontaneous movement begins. In *Drosophila melanogaster* brief muscle twitches start at 840 minutes during embryogenesis, and become more frequent as development progresses, while on the 1020 minutes stage a burst of activity in muscles is shown (Crisp et al. 2008). In *Danio rerio* spontaneously-occurring motor output occurs between 18 and 27 hours post fertilization (Menelaou et al. 2008). Interestingly, although in *C. elegans* we found clearly elevated mtDNA gene expression with only few members of the nuclear OXPHOS, such elevation started around the beginning of muscle movement, namely after ∼430 minutes (quickening stage) (Chisholm and Hardin 2005). Hence, elevation of OXPHOS gene expression - both mtDNA and nuclear DNA-encoded OXPHOS genes - precedes locomotion-associated developmental stages in the frog, zebrafish, drosophila and possibly in the worm. Association between upregulation of mito-nuclear gene expression and the initiation of locomotion is especially attractive since many mitochondrial disorders display impaired movement phenotypes (Tranchant and Anheim 2016; Ticci et al. 2021), suggesting the importance of CMNGE for such disorders – an interpretation that should be tested in future experiments.

Our results strongly support a conserved regulatory basis underlying CMNGE during the transition from fetal to neonatal life in many tissues. Since this conclusion is based on gene expression analysis at the RNA level, it would be interesting to test for mito-nuclear gene expression coordination during development at the protein and translation levels. That would be of great interest in light of the claim for strong impact of mitochondrial translation on the co-regulation of the two genomes (McShane et al. 2024). With this in mind, it would be of interest to explore additional tissues, particularly those with lower versus higher bioenergetic activity, and to conduct a comparative analysis of CMNGE at tissue-specific, cell type levels, as well as at the RNA and protein levels. This should be performed in the future once relevant experimental datasets become available.

In summary, our study revealed coordination of gene expression between mitochondria and the nucleus in the transition from fetal to neonatal life, and for the first time identified its conservation in vertebrates. Unexpectedly, our analysis revealed elevated CMNGE which reflected a conserved metabolic shift post-birth (mammals) or hatching (bird and lizard). Key regulatory factors, e.g. PGC-1α, NRF2, TFB2M and C/EBPB emerge as potential regulators of such CMNGE shifts. Secondly, we found that elevated CMNGE preceded the initiation of locomotion in frog, zebrafish, and drosophila. Our findings pave the path towards future identification of the underlying mechanism of CMNGE during development, as well as in particular tissues, during the course of evolution.

## Methods

### Data acquisition

Data source details and number of samples per analyzed dataset are indicated in Table 1.

For Dataset I, raw bulk RNA-seq fastq files were downloaded from ArrayExpress (https://www.ebi.ac.uk/arrayexpress/) with the following accession codes: E-MTAB-6769 (chicken), E-MTAB-6782 (rabbit) E-MTAB-6798 (mouse), E-MTAB-6811 (rat), E-MTAB-6813 (rhesus), E-MTAB-6814 (human) (Cardoso-Moreira et al. 2019). For data analysis – see RNA-seq analysis section. Dataset 1 included 1,346 RNA-seq libraries, encompassing embryo, fetal and neonatal development in seven organs, 9-23 developmental stages. Samples that were collected during the same developmental week were considered biological replicates. For instance, samples that were generated from Carnegie stages 13 and 14 were considered as replicates (4 wpc), as previously performed (Cardoso-Moreira et al. 2019). Accordingly, the data included 1-4 replicates per stage. Briefly, human prenatal samples were available at 4 weeks post-conception (wpc) and at each week until 20 wpc (except for 14, 15 and 17 wpc). Notably, there were no human samples available between 20 and 38 wpc. Postnatal human samples included ‘infants’ (6-9 months old), “toddlers” (2-4 years old), “school” (7-9 years old), “teenagers” (13-19 years old) and adults from each decade until 63 years of age. Human ovary samples were collected only from prenatal samples (until 18wpc) and kidney development was sampled up until 8 years of age (“school”). Samples from rhesus monkey (*Macaca mulatta*) were collected while taking into account that gestation lasts ∼167 days. Rhesus samples were collected from fetal stage e93 and from five stages prior to birth (until e130). Notably, ovary was not sampled from rhesus. For mouse (*Mus musculus*) samples were collected from day e10.5 and during subsequent days, each day, until birth (i.e., until e18.5). Postnatal mouse samples included P0, P3, P14, P28 and P63. For rabbit - *Oryctolagus cuniculus*, outbred New Zealand breed, samples were collected starting from e12 and subsequent 11 time points up until (including) e27, while taking into account that gestation length is 29-32 days. Postnatal rabbit samples included P0, P14, P84 and between P186-P548. For rat - *Rattus norvegicus*, outbred strain Holtzman SD, the time-series start at e11 and include daily time points until birth (until e20). Postnatal samples include P0, P3, P7, P14, P42 and P112. For chicken - *Gallus gallus*, red junglefowl, egg incubation lasts ∼21 days. Thus, collection time series started at e10 while sampling three additional stages until e17. Postnatal samples include P0, P7, P35, P70 and P155.

For Dataset II, normalized count matrix of processed transcripts per million (TPM) was downloaded from GEO accession number GSE176387. We analyzed five developmental stages from porcine liver: embryonic days E38, E80, the day of birth (0d); weaning at 28 days (28D); sexual maturity at 110 days (110D), with 6 replicates in each stage.

For Dataset III, FPKM (Fragments Per Kilobase Million) normalized count matrix was downloaded from GEO accession number GSE121019. We analyzed three developmental stages available for chicken liver: embryonic day of 13, postnatal stages at 5 weeks and 42 weeks of age, with each having three replicates.

For Dataset IV, raw read count matrix was downloaded from GEO accession number GSE58455. We analyzed seven independently collected E13.5 samples and 12 adult samples (6 at 6 weeks of age, 6 at 16 weeks of age) from mouse heart.

For Dataset V (*Anolis carolinensis*) we took into account that Annolis development typically lasts 30-33 days and include 19 stages (Sanger et al. 2008). Raw bulk RNA-seq fastq files were downloaded from GEO accession number GSE97367. We analyzed 22 embryonic samples (from stages 15-16) and 30 adult samples from four organs (brain, heart, kidney, liver) and twenty-six whole embryonic samples.

For Dataset VI (*Xenopus tropicalis*) raw data fastq files were downloaded from GEO accession number GSE37452. The samples included two clutches: A. From two-cells to stage 45, and (B) from stage 9 to stage 42. These stages span from the beginning of cleavage to the tailbud free swimming stage. In all cases whole organism was sampled.

For Dataset VII, raw reads count matrices for the zebrafish (*Danio rerio*), *Drosophila melanogaster* and *C. elegans* were downloaded from GEO accession number GSE70185 (Levin et al. 2016). Time series included samples from the cleavage stage up to the time point prior to hatching.

### RNA-seq analyses: mapping, read count and control for nuclear mitochondrial sequences - NUMTs

For Datasets I, V and VI, sequenced reads were trimmed using Trim Galore (version 0.4.5; https://www.bioinformatics.babraham.ac.uk/projects/trim_galore/). Prior to reads count analyses, we controlled for possible noise originating from nuclear mitochondrial fragments (NUMTs), e.g. mtDNA sequence fragments that were transferred from the mitochondria to the nucleus during the course of time (Mishmar et al. 2004). NUMTs potentially pose an obstacle to mtDNA gene expression assessment, as a subset of RNA reads, particularly those that are relatively recent (Mishmar et al. 2004), might be more similar to the active mtDNA and sequencing reads from mtDNA genes; such fragments might be erroneously filtered out while applying the unique mapping protocol. To overcome such problem, reads were firstly mapped solely against the mtDNA genome using bwa using the aln parameter (BWA-backtrack algorithm) (Li and Durbin 2009); then, the reads that did not map to the mtDNA, were uniquely mapped against the entire reference genome of the relevant organisms using STAR (version 2.5.3), while employing default parameters, and the parameter [—outFilterMultimapNmax 1] (Dobin et al. 2013). Expression levels of all genes were counted using HTSeq-count v0.11.2 (Anders et al. 2015), using default parameters while employing the [-f bam] parameters. High quality samples were used as listed in the original study (Cardoso-Moreira et al. 2019).

Libraries with non-zero read counts of mtDNA genes were used for subsequent analysis to avoid technical noise. Reference genomes that were used are listed in Table S2.

### Quantification of gene expression and statistical analysis

Read counts data matrices were normalized using DESeq normalization method available from the DESeq2 R package (Love et al. 2014). For Dataset I, the statistical analysis we used a median of 2-4 replicates for each stage. Heatmaps were created using the pheatmap R package (Kolde 2012). To identify the developmental time point presenting with the highest fold-change (FC) of mtDNA gene expression, while considering the median of replicates from each stage, sliding window algorithm was employed with length of w=2 (two consecutive samples), w=4 (four consecutive samples). For a larger dataset (Dataset VII, having more than N=55 per analysis), we used a window of w=16. Log2FC values were calculated for each time point in each window. To detect the largest change in mtDNA gene expression in more than one time point, we used a window of four time points. Differential expression of mitochondrial genes in a given time point was calculated (DESeq2) separately for each dataset, while considering samples from the previous and subsequent time points. Multiple testing correction for each specified set of genes was performed using adjusted p-value value of 0.05. Box plot and bar plot graphs were generated using “ggplot2” (Wickham and Wickham 2016) and RColorBrewer R packages. Tables were generated using the R packages ‘openxlsx’, ‘tidyr’ and ‘reshape2‘ (Zhang 2016). Level of coordination of mito-nuclear genes was calculated as follows: the number of significantly upregulated OXPHOS genes (*X*_*up*_) minus the number of significantly downregulated OXPHOS genes (*X*_*down*_), divided by the number of total available OXPHOS genes, as in the following formulae:

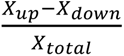

Expression of genes was considered significantly altered if their adjusted p-value was smaller than 0.05 and the absolute value of log2FC was greater than 0.2.

### GO terms used in the analysis

The following Gene Ontology (GO) Terms were used to download and perform differential expression analysis: Genes belonging to ‘TCA’ (GO:0006099), ‘glycolytic process’ (GO:0006096), ROS formation (GO:1903409), ‘Locomotion’ (GO:0040011), and ‘positive regulation of cellular response to hypoxia’ (GO:1900039). We assessed the expression of candidate mtDNA regulatory factors of transcription, regulatory factors of mtDNA replication, and nuclear DNA-encoded factors with known mitochondrial RNA-binding activity (Wolf and Mootha 2014; Cohen et al. 2016), in addition to RNA and DNA-binding proteins that were recently identified in human mitochondria (Ardail et al. 1993; Fernández-Vizarra et al. 2008; She et al. 2011; Blumberg et al. 2014b; Lambertini et al. 2015; Chatterjee et al. 2016). Hypoxia gene set was downloaded from a previous study (Bono and Hirota 2020). The top 100 responding genes with the highest score in humans were used for the analyses.

## Acknowledgements.

This work was funded by grants from the Israeli Science Foundation (ISF 404/21) and from the life sciences division of the US Army Research Office (grant number 80581-BB) awarded to DM. The authors wish to thank Ben-Gurion University for awarding HM the Negev scholarship for excellent PhD students and a short-term Postdoc training grant.

